# Gland-Mass Adaptation Explains the Cortisol Paradox in PTSD

**DOI:** 10.1101/2025.10.13.681561

**Authors:** Dor Danan, Yaniv Grosskopf, Yoav Hayut, Yoel Toledano, Keren Doenyas-Barak, Avi Mayo, Uri Alon

## Abstract

Post-traumatic stress disorder (PTSD) often shows an endocrine paradox: basal cortisol is low while ACTH, its upstream driver, is normal. Hormone kinetics alone cannot produce this: at steady state the two move together. We show that slow adaptation of pituitary and adrenal functional mass, in a previously calibrated model, resolves the paradox: gland mass acts as an integral feedback that holds ACTH at its set point while cortisol falls. Only two changes can lower basal cortisol, reduced central drive or increased glucocorticoid receptor (GR) sensitivity, and they move GR occupancy in opposite directions. A twofold increase in GR sensitivity, consistent with dexamethasone suppression tests, reproduces low cortisol with normal ACTH, lower cortisol during acute stress and a blunted dexamethasone–CRH response. Cortisol in PTSD is thus not too low, but too high for the receptors that read it. In records from Israel’s largest health provider, morning cortisol is 4.5% lower in PTSD across 180,000 tests, and eight of nine glucocorticoid-responsive markers shift toward glucocorticoid excess, the sign only body-wide sensitization predicts.

## 1 Introduction

Relative hypocortisolemia in PTSD has resisted mechanistic explanation for three decades (Yehuda et al., 1990). Meta-analyses across plasma, urinary and salivary measures support reduced basal cortisol, though with substantial heterogeneity attributable to sampling conditions, comorbidity and diagnostic breadth (Meewisse et al., 2007; Pan et al., 2018, 2020). Reduced basal corticosterone is likewise reported in many rodent PTSD paradigms (Algamal et al., 2018; Tseilikman et al., 2021; Verbitsky et al., 2020).

Low basal cortisol on its own would be easy to explain, since many perturbations can produce it. What resists explanation is its pairing with normal ACTH (Cooper et al., 2017; Golier et al., 2012; Kanter et al., 2001; Yehuda et al., 2004a). Because ACTH drives cortisol, the two normally rise and fall together. Low cortisol with normal ACTH means something other than hormone kinetics sets their relationship.

Enhanced glucocorticoid receptor (GR) sensitivity is the standard candidate mechanism. PTSD is associated with heightened GR sensitivity (Rohleder et al., 2004, 2010; Somvanshi et al., 2020), increased GR expression (Labonté et al., 2014; van Zuiden et al., 2012; Yehuda et al., 1991), and stronger negative feedback (de Kloet et al., 2006; McFarlane et al., 2011; Yehuda et al., 1993, 2004a, 2014). Stronger feedback alone, however, predicts suppression of ACTH along with cortisol, which is not what is observed (Lawrence and Scofield, 2024; Yehuda, 2009; Zoladz and Diamond, 2013). Existing models capture parts of the phenotype, linking GR sensitivity to inflammation (Somvanshi et al., 2020), classifying cortisol time series (Sriram et al., 2012), or generating low cortisol as an alternative stable state under bistability (Kim et al., 2016, 2018), but all treat secretory capacity as fixed and so inherit the same constraint.

Endocrine glands change functional mass over weeks in response to their trophic inputs: ACTH enlarges the adrenal cortex (Lotfi and De Mendonca, 2016; Swann, 1940; Ulrich-Lai et al., 2006), CRH expands pituitary corticotrophs (Asa et al., 1992; Bruhn et al., 1984; Westlund et al., 1985), and prolonged glucocorticoid excess shrinks them (Beuschlein et al., 2024; Nolan and Levy, 2001; Roussel-Gervais et al., 2016). A previously calibrated model incorporating these dynamics accounts for months-scale hormonal imbalance after prolonged stress (Karin et al., 2020), an opponent process in alcohol addiction (Karin et al., 2021), seasonal variation (Tendler et al., 2021), and the failure of most HPA-targeted drugs in chronic stress (Milo et al., 2025).

Here we show that gland-mass adaptation reproduces the endocrine phenotype of PTSD with a single parameter change, a twofold increase in GR sensitivity consistent with in-vivo feedback tests, and that it constrains which changes in the axis are observable at all. Adaptation pins ACTH and leaves basal hormone levels insensitive to most axis parameters, narrows the changes that can lower cortisol to two, and supplies a signed test that separates them: glucocorticoid action in peripheral tissue. We use population cortisol to set the absolute operating point of the axis, and carry out the signed test in routine laboratory markers from the same records.

## 2 Results and Discussion

### 2.1 Increased GR sensitivity in the gland-mass model gives low cortisol with normal ACTH

The model tracks CRH (*x*_1_), ACTH (*x*_2_), cortisol (*x*_3_), and the functional masses of pituitary corticotrophs (*P*) and adrenal cortex (*A*) (Fig. 1; Methods). Cortisol feeds back through mineralocorticoid receptors (MR) in the hypothalamus and through GR at both hypothalamus and pituitary, with GR occupancy a Hill function of *x*_3_*/K*_GR_, where *K*_GR_ is the half-maximal parameter; a smaller *K*_GR_ means a more sensitive receptor, and we use GR sensitivity for 1*/K*_GR_. The masses grow in proportion to their trophic drive and turn over at a fixed rate.

**Figure 1:**
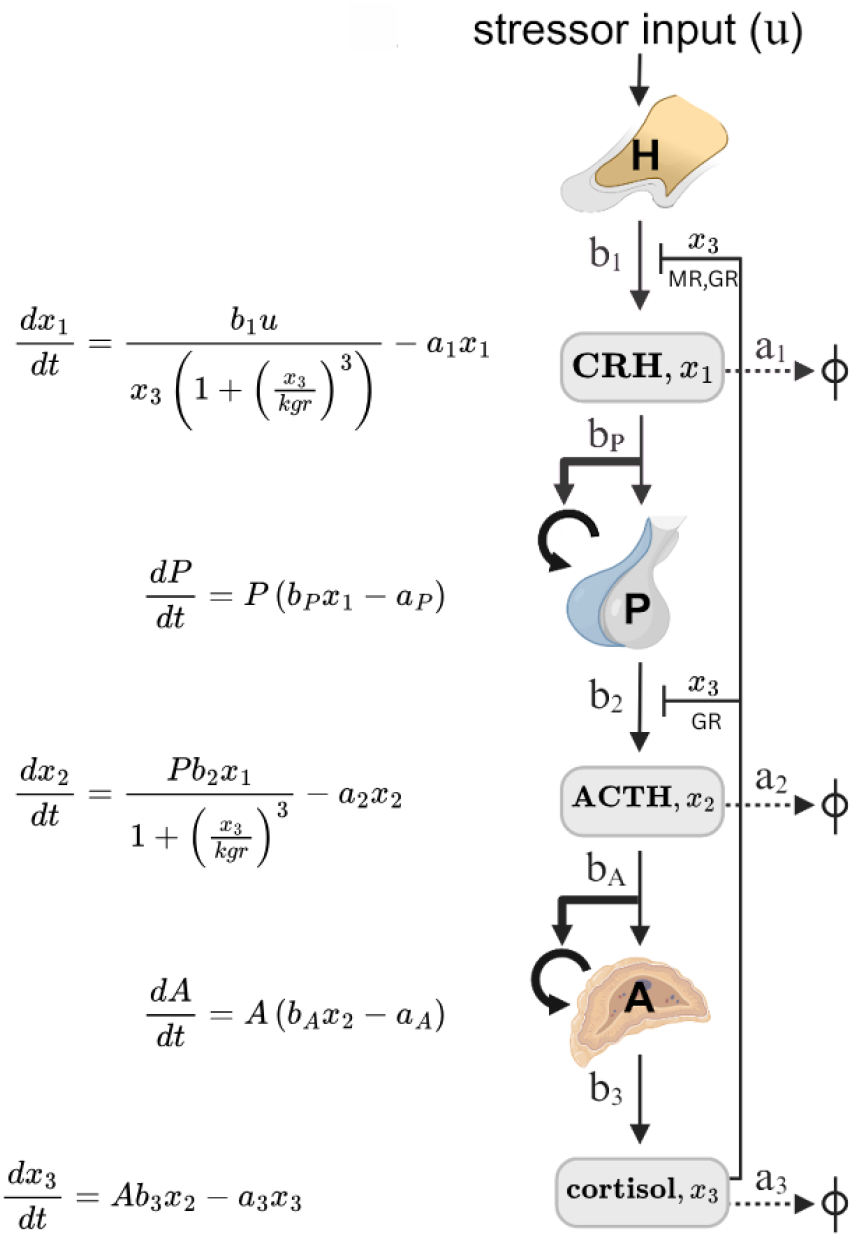
The HPA circuit diagram and corresponding equations. The hypothalamus H secretes corticotropin-releasing hormone (CRH) at a rate *b*_1_ in response to a stressor input *u*. CRH causes the pituitary (P) to secrete adrenocorticotropic hormone (ACTH) at a rate *b*_2_ and to grow in functional mass at a rate *b_P_*. ACTH signals the adrenal gland (A) to secrete cortisol at rate *b*_3_ and to grow in functional mass at rate *b_A_*. Cortisol inhibits ACTH secretion at the pituitary through glucocorticoid receptors (GR), and CRH secretion at the hypothalamus through both GR and mineralocorticoid receptors (MR). The hormone removal rates are *a*_1_, *a*_2_ and *a*_3_ for CRH, ACTH and cortisol. Thick arrows indicate the interactions that affect gland sizes on the scale of months. The corresponding equation is given to the left of each interaction.

PTSD is modeled as a twofold increase in GR sensitivity (Danan et al., 2021; Matić et al., 2015; Yehuda et al., 2004b): *K*_GR_ = 2.5 for PTSD and *K*_GR_ = 5 for controls, in units normalized to baseline cortisol. The twofold ratio is consistent with in-vivo feedback tests, and the absolute scale is set by population cortisol (both below). Every other parameter follows the previously published calibration (Karin et al., 2020), except cortisol clearance, which is raised to *a*_3_ = 0.06 min^−1^ to match free-cortisol disposition kinetics (Dorin et al., 2022) (Appendix Table S1; Methods).

At *K*_GR_ = 2.5, steady-state cortisol is 4.4% below the control value, ACTH is unchanged, adrenal mass is 4.4% lower and corticotroph mass 4.6% higher (Fig. 2). Reduced adrenal mass is documented in rat PTSD models (Manukhina et al., 2018; Tseilikman et al., 2021). The predicted corticotroph change corresponds to roughly 4 mm^3^ given a pituitary volume of ∼450 mm^3^ of which ∼20% is corticotrophs (Berntsen et al., 2021; Melmed et al., 2022), which lies well below inter-individual variance and is consistent with the conflicting imaging reports in PTSD (Atmaca et al., 2017; Cooper et al., 2017; Thomas and De Bellis, 2004). The invariance of ACTH is forced by the mass equations, as the next section shows.

**Figure 2:**
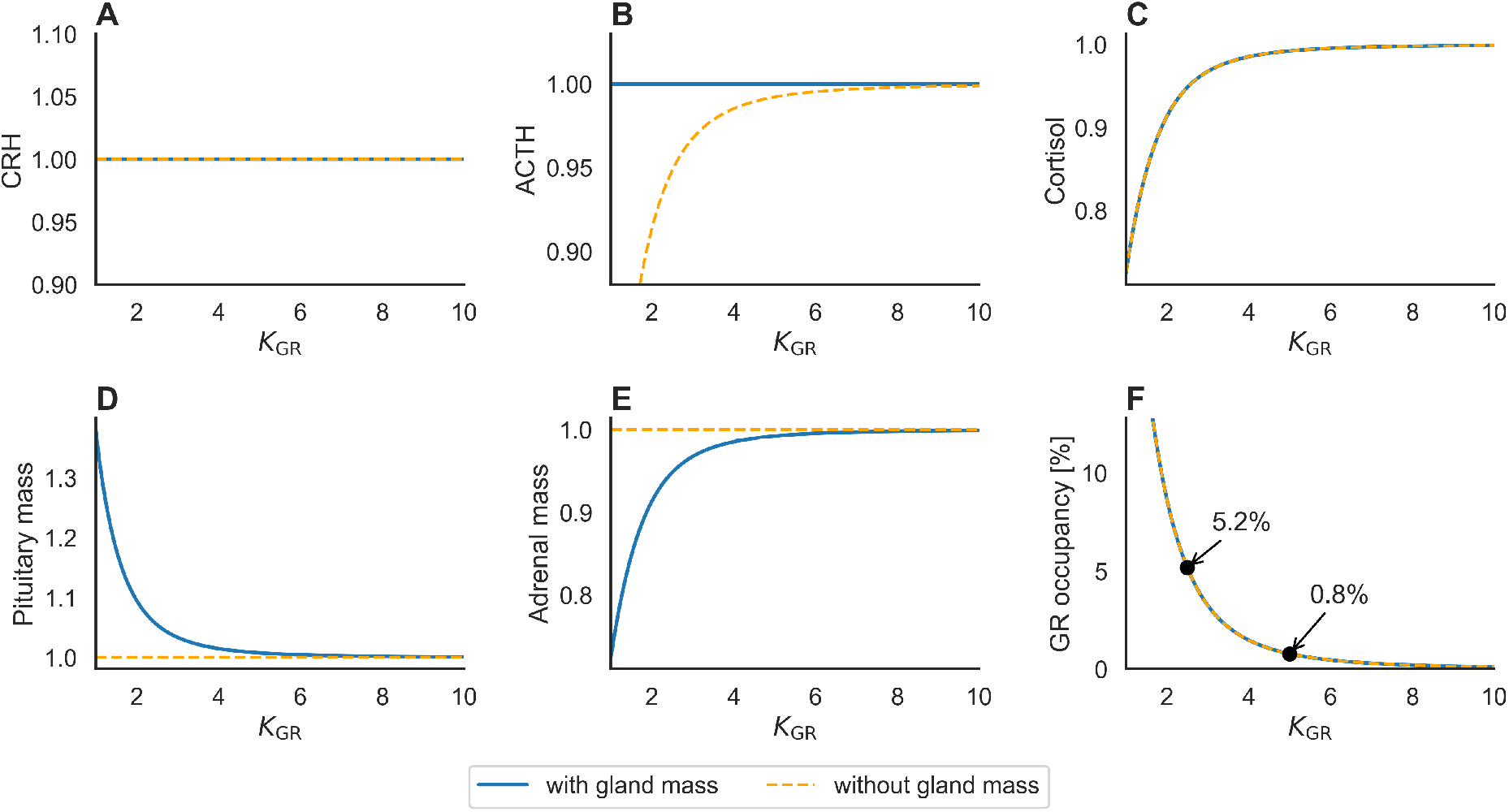
The steady-state phenotype across GR sensitivity, with and without gland-mass adaptation. Each panel is one model variable at steady state as a function of *K*_GR_: (A) CRH, (B) ACTH, (C) cortisol, (D) pituitary corticotroph mass, (E) adrenal cortex mass, (F) fractional GR occupancy. Blue, the full model with adaptive masses; dashed orange, the same model with the masses clamped to 1. The two coincide for CRH, cortisol and GR occupancy and diverge for ACTH and the masses: only with adaptation does ACTH hold at baseline as *K*_GR_ falls (B), which is the PTSD pattern of low cortisol with normal ACTH. Occupancy, (*x*_3_*/K*_GR_)*^n^/*[1 + (*x*_3_*/K*_GR_)*^n^*] with *n* = 3 evaluated at the steady-state cortisol, is annotated at the PTSD (*K*_GR_ = 2.5) and control (*K*_GR_ = 5) operating points: it rises from 0.78% to 5.2%, an increase of 4.4 percentage points reached at lower circulating cortisol, with the receptor far below saturation in both groups.

### 2.2 Gland-mass adaptation pins ACTH and makes most of the HPA axis unobservable

The mass equations constrain the steady state for any parameter values. A gland grows when its trophic input is above a threshold and shrinks when it is below, so each mass equation is an integral feedback loop (Karin et al., 2016), and such a loop at rest has zero error: at steady state the input must sit exactly at the threshold, whatever else in the axis has changed. CRH and ACTH therefore show exact adaptation. Setting *dP/dt* = *dA/dt* = 0 for non-zero masses gives

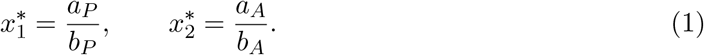

CRH and ACTH are fixed by the ratio of each gland’s turnover to its trophic gain. Neither depends on *K*_GR_, on the strength of glucocorticoid feedback, or on anything downstream. ACTH is therefore not a fitted output, and cannot move in response to a change in GR sensitivity (Fig. 2B). Cortisol is free to fall, because adrenal mass absorbs the difference: substituting Eq. (1) into the hormone steady states gives *A^∗^* ∝ *x*_3_ and *P^∗^* ∝ 1/*x*_3_. A smaller adrenal cortex delivers less cortisol per unit ACTH, so the axis reaches a new steady state with lower cortisol at an unchanged ACTH set point.

The pinning has a consequence beyond ACTH. Writing out the remaining steady-state conditions gives

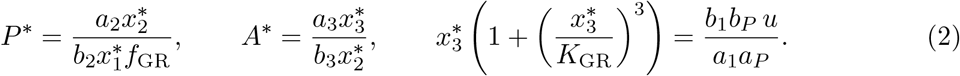

The pituitary and adrenal secretion and removal constants *b*_2_*, a*_2_*, b*_3_*, a*_3_ appear in the mass expressions and nowhere else, and are absent from all three hormone levels. A change in adrenal output per unit ACTH, or in corticotroph output per unit CRH, therefore leaves basal hormone levels unchanged: the gland resizes and the hormone returns to where it was. Numerically, a 20% change in any of the four does not change CRH, ACTH or cortisol at all; only the corresponding gland mass moves. That gland-mass feedback absorbs changes in secretory parameters is the dynamical compensation described by Karin et al. (2016). Its consequence for inference is that in the HPA axis only two parameter combinations remain visible in basal hormones, and they move GR occupancy in opposite directions (next section), which turns a peripheral readout of glucocorticoid action into a signed test between them.

Without adaptation these constraints vanish. With *P* and *A* clamped, *x*_2_ ∝ *x*_3_, ACTH falls with cortisol (Fig. 2B, dashed), and every secretory parameter becomes visible again. The two model variants agree exactly on CRH, cortisol and GR occupancy and disagree only on ACTH and the gland masses (Fig. 2), which isolates gland-mass adaptation as the necessary ingredient. The low-cortisol, normal-ACTH signature is thus a generic property of an endocrine axis whose secretory capacity adapts, rather than evidence of an unusual perturbation, and in such an axis most perturbations leave no trace at all.

Because the pinning is an equilibrium property, it also predicts a transient. Stepping *K*_GR_ from 5 to 2.5 at *t* = 0, cortisol reaches its new level within hours and ACTH falls with it by 4.4%, the fixed-capacity behavior. ACTH is then restored as the glands re-adapt, recovering to −1.3% by 30 days and to baseline by 60, on the timescale set by corticotroph and adrenal turnover (10 and 20 days). A fixed-capacity axis leaves that deficit in place permanently. The normal ACTH of chronic PTSD is therefore a property of an axis observed long after onset, which is the regime the cross-sectional comparison below samples.

### 2.3 Only two changes can lower basal cortisol, and they move GR occupancy in opposite directions

Equation (2) makes basal cortisol depend on the model’s parameters only through two combinartions: the effective central drive *b*_1_*b_P_ u/*(*a*_1_*a_P_*) and *K*_GR_ (with the Hill exponent fixed). Every other parameter, including cortisol clearance *a*_3_, is absorbed by gland mass.

Of the two changes, reduced drive is the less plausible: PTSD involves heightened rather than reduced stress exposure and arousal. Higher drive alone would raise cortisol, so the observed fall points to increased GR sensitivity. The two changes can also be told apart directly. Tuned to the same 4.4% fall in cortisol, reduced drive lowers GR occupancy at the feedback nodes from 0.78% to 0.68%, whereas raised sensitivity takes it to 5.2%: identical in hormone levels, opposite in the quantity that acts on tissue. A third account, sensitization confined to the feedback loop, behaves like reduced drive in the periphery, where unchanged receptors see 4.4% less cortisol and occupancy also falls to 0.68%. Only body-wide sensitization raises peripheral occupancy, so a peripheral readout of glucocorticoid action rules out the other two with a single sign.

Adaptation also localizes the change. Allowing *K*_GR_ to differ between the hypothalamic and pituitary arms, pituitary sensitization alone moves no hormone and is absorbed into a 5.4% larger corticotroph mass, whereas hypothalamic sensitization alone reproduces the full cortisol fall. The endocrine phenotype of PTSD is therefore evidence about the hypothalamic feedback arm, and dexamethasone, which acts at the pituitary on a timescale too short for mass adaptation, is the complementary probe.

### 2.4 Enhanced GR feedback reproduces dexamethasone hypersuppression, lower cortisol during acute stress and a blunted Dex–CRH response

The twofold ratio is consistent with the dexamethasone suppression test, the most direct in-vivo read of glucocorticoid feedback. With the dexamethasone strength set so that the control arm matches the observed 72–78% suppression, the PTSD arm reaches 88–91%, a 2.3-fold lower residual cortisol, against 2.4-fold in PTSD without comorbid depression (Yehuda et al., 2002) and 1.5- to 3.5-fold across studies (Newport et al., 2004; Yehuda et al., 2002) (Methods); ACTH is likewise more suppressed after dexamethasone in PTSD (Yehuda et al., 2004a). Dexamethasone acts mainly at the pituitary, so this tests the pituitary arm; applying the same ratio to the basal phenotype, which the hypothalamic arm sets (above), assumes a single sensitization at both sites, an assumption the peripheral markers below support. Leukocyte assays show smaller differences in dexamethasone IC_50_, 1.2–1.6-fold (Blalock et al., 2024; Rohleder et al., 2004; Yehuda et al., 2004b), and some show none or the opposite (de Kloet et al., 2007; Matić et al., 2013).

The same parameter change reproduces both standard dynamic challenges without refitting any feedback parameter. At low *K*_GR_, a 20-minute stress pulse gives lower cortisol throughout the test with a smaller rise, as in salivary cortisol during the Trier Social Stress Test (Fig. 3A) (Metz et al., 2020; von Majewski et al., 2023; Zorn et al., 2017). A dexamethasone–CRH protocol produces a lower ACTH baseline and a ∼3-fold lower plateau after exogenous CRH, matching Ströhle et al. (2008) (Fig. 3B). Two opposing influences act during that challenge: enhanced GR feedback suppresses ACTH secretion, while the larger corticotroph mass raises secretory capacity and partly offsets it, and the net effect is the blunted response observed.

**Figure 3:**
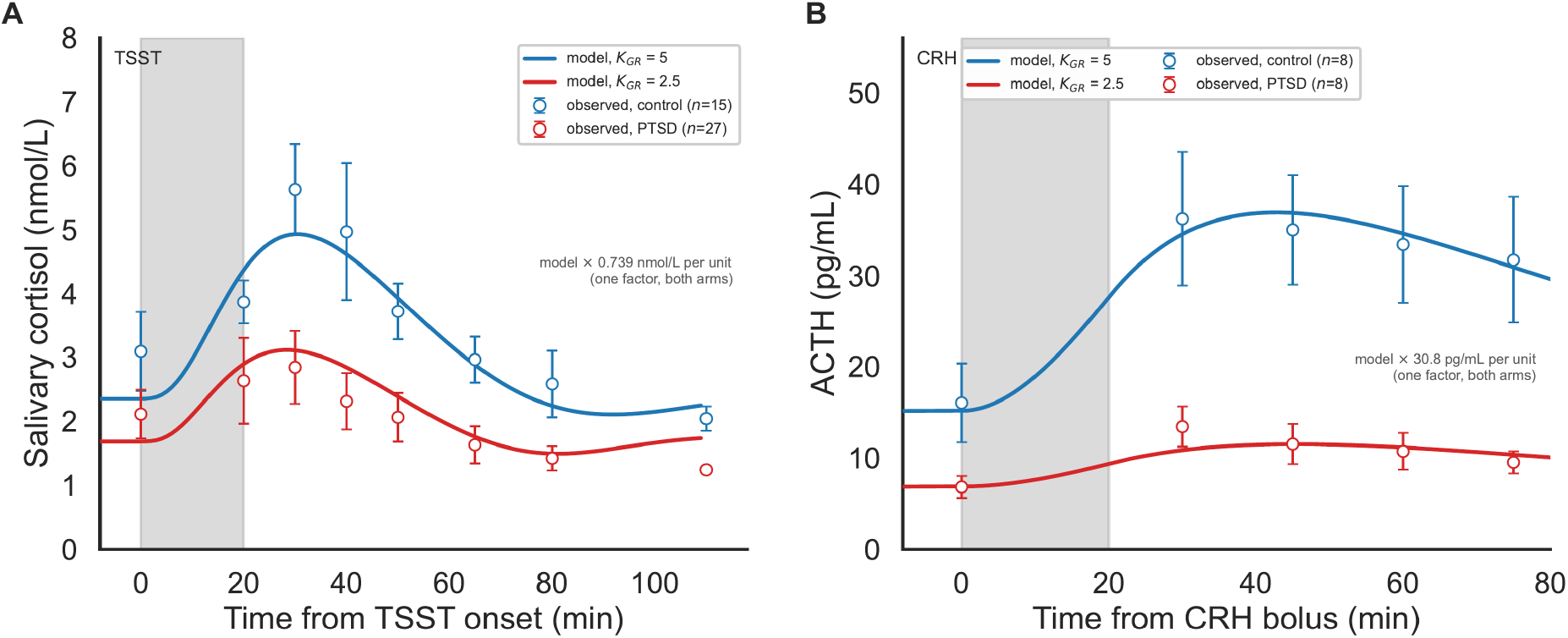
The same parameter change reproduces both standard endocrine challenges. Measurements (open symbols, mean ± SEM) are shown with the model (lines) on the same axes; red, PTSD (*K*_GR_ = 2.5); blue, control (*K*_GR_ = 5). No feedback parameter was refitted to either dataset; the TSST pre-stress drive and pulse size and the two Dex–CRH input magnitudes, each shared by both arms, were fitted (Methods). (A) Salivary cortisol during the Trier Social Stress Test, a 20-minute stress pulse (gray band) simulated as a tenfold increase in the stress input *u*; data from von Majewski et al. (2023) (*n* = 27 PTSD, *n* = 15 controls). The low-*K*_GR_ axis has lower cortisol throughout, with a smaller rise. (B) ACTH in the dexamethasone–CRH test, with dexamethasone as a constant cortisol-equivalent GR agonist and a 20-minute exogenous CRH pulse from *t* = 0 (gray band); data from Ströhle et al. (2008) (*n* = 8 per arm; 0.75 mg dexamethasone at 23:00 and 100 *µ*g CRH at 15:00). The low-*K*_GR_ axis has both a lower Dex-suppressed level and a lower plateau. The model is normalized, so each panel carries one display factor converting model units to the measured units, fitted by least squares and identical for the two arms (Methods); the measurements were digitized from the published panels. The TSST run has a pre-stress *u* = 4, the Dex–CRH run *u* = 1 (Methods).

The TSST data are best fitted with a pre-stress drive of *u* = 4, fourfold above the operating point used elsewhere, and the model then under-predicts the measured peak reduction (Methods).

Other challenge studies report enhanced ACTH responses to CRH in PTSD (Rasmusson et al., 2001), reviewed in de Kloet et al. (2006), and the group sizes throughout are small (*n* = 8 per arm in Ströhle et al., 2008). The simulations correspond to the blunted pole of this literature.

### 2.5 The model predicts elevated GR occupancy despite low cortisol

Because GR is far from saturation at baseline, the Hill term can be expanded: fractional occupancy is ≈ (*x*_3_*/K*_GR_)^3^ when that ratio is small, so occupancy is governed by the cortisol-to-sensitivity ratio. That ratio is 0.198 at the control operating point and 0.379 at the PTSD operating point, and occupancy at the feedback nodes accordingly goes from 0.78% to 5.2%, an increase of 4.4 percentage points (Fig. 2F).

Moderate Cushing’s syndrome, threefold cortisol acting on a normal receptor, raises occupancy to 17.4%, an increase of 16.6 percentage points. The PTSD change is therefore about a quarter of that of moderate hypercortisolism in the quantity tissue responds to, and is reached at lower circulating cortisol. Sustained GR activation is associated with hippocampal dendritic retraction, impaired neurogenesis and volume loss (Brown et al., 2015; Conrad, 2008; Kino, 2015; McEwen et al., 2016).

The occupancy figures depend on the assumed cooperativity: *n* = 2 and *n* = 4 give PTSD occupancies of 11.2% and 2.3%, so the increase spans 2.1 to 7.6 percentage points against 4.4 at *n* = 3; Fig. 4C shows the corresponding predicted cortisol reductions. The dependence on the operating point is of similar size (below).

**Figure 4:**
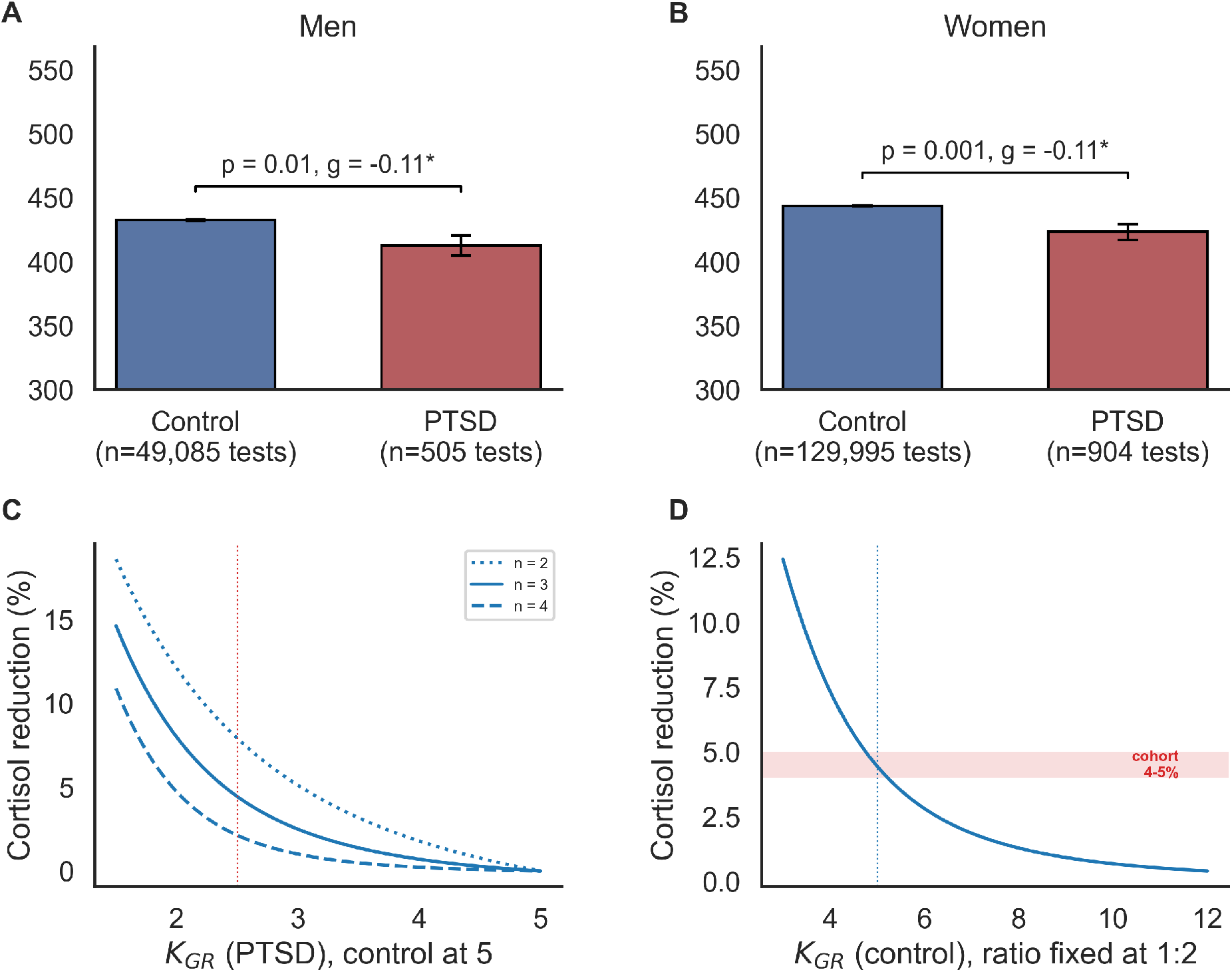
Population cortisol constrains where the axis operates. (A, B) Directly age-standardized morning serum cortisol (nmol/L, 06:00–10:00, ages 18–70) in the Clalit cohort for men and women; both groups are standardized to the combined age distribution, so the bars differ only by the PTSD effect. Error bars are the standard error of the standardized mean, propagated from the per-bin values, so the small case groups govern the uncertainty; *n* is the number of cortisol tests. Tests within 90 days after a corticosteroid dispensing are excluded, and PTSD tests are those after the first PTSD diagnosis (Methods). (C) Predicted cortisol reduction as the PTSD *K*_GR_ is swept with the control held at 5, for Hill coefficients *n* = 2, 3, 4; the dotted vertical marks *K*_GR_ = 2.5. The predicted reduction depends on the assumed cooperativity, as does the resulting change in GR occupancy. (D) Predicted reduction as the absolute scale is slid with the PTSD-to-control *K*_GR_ ratio held at 1:2; the shaded band spans the observed 4–5%, which selects a control *K*_GR_ of ≈ 5 (dotted vertical at 5).

### 2.6 Morning serum cortisol is about 4.5% lower in PTSD and constrains where the axis operates

We compared morning serum cortisol between adults with and without a PTSD diagnosis in the Clalit health-maintenance organization, Israel’s largest, with over five million members (Balicer and Afek, 2017; Bar et al., 2025; Cohen et al., 2021). After restricting to 06:00–10:00 draws, excluding diagnoses known to perturb the axis, excluding tests drawn within 90 days after a corticosteroid dispensing, using only PTSD tests taken after the diagnosis, and directly age-standardizing, cortisol was lower in PTSD by a similar amount in both sexes: 4.6% in men (Δ = −19.8 nmol/L, *g* = −0.11, *p* = 0.012; 404 cases, 38,085 controls) and 4.5% in women (Δ = −20.1 nmol/L, *g* = −0.11, *p* = 0.001; 730 cases, 97,814 controls) (Fig. 4A,B; Appendix Table S2). On the log scale the geometric-mean reduction is 5.1% in men and 4.5% in women, and per-bin medians give 7.3% and 4.7%.

The feedback tests fix the PTSD-to-control ratio of *K*_GR_, not its absolute scale. With the PTSD-to-control ratio held at 1:2, the predicted cortisol reduction runs from 1.3% (PTSD/control *K*_GR_ = 4*/*8) to 20.7% (1*/*2), and the observed ≈4.5% selects a control *K*_GR_ of ≈ 5 in units of baseline cortisol (Fig. 4D). With baseline free cortisol of ≈10 nM this is ≈50 nM, the in-vitro EC_50_ of cortisol at GR (35–67 nM; Levitt, 2024). Over admissible reductions of 2–10%, the control *K*_GR_ moves from 6.8 to 3.4 while the predicted rise in GR occupancy stays positive (2.0–9.8 percentage points).

### 2.7 Peripheral laboratory markers show elevated glucocorticoid action in PTSD

The signed test derived above can be carried out in routine blood chemistry: reduced drive and feedback-confined sensitization predict lower peripheral glucocorticoid action, body-wide sensitization higher.

Nine routinely measured blood analytes have documented responses to glucocorticoid excess. Before any group comparison, each was assigned the direction it moves under glucocorticoid excess, taken from the pharmacological literature: white blood cells (Bauer et al., 2018; Crockard et al., 1998), platelets (Isidori et al., 2015), hemoglobin (Varricchio et al., 2023), glucose (Cho and Suh, 2024), HbA1c (Herndon et al., 2021) and triglycerides (Taskinen et al., 1983) rise, while C-reactive protein (CRP) (Michel et al., 2006; Vogel et al., 2023), potassium (Quinkler and Stewart, 2003) and TSH (Cavalieri, 1991) fall. We then asked whether each marker moves in its assigned direction in 40,240 PTSD cases against 311,362 controls from the same records, matched on sex, birth year and primary-care clinic, in five windows around the diagnosis date (for a control, the diagnosis age of its matched case; Methods): baseline (−10 to −6 y), prodrome (−6 to −1 y), diagnostic (−1 to +0.5 y), early (+0.5 to +3 y) and late (+3 to +10 y) (Fig. 5B; Methods). We report concordance as *k/m*: of the *m* markers that pass a 5% false-discovery threshold, *k* move in the assigned direction. In the prodrome eight of the nine markers pass and the result is 8/8 (sign test *p* = 0.004; Fig. 5A); in the early window it is 8/8, in the late window 7/7, and at baseline 6/7 (*p* = 0.06). The concordance holds separately in men (prodrome 6/6, diagnostic 6/9, early 5/5, late 5/5) and in women (8/9, 6/7, 6/6 and 7/7 in the same order), with the discordance in both sexes concentrating in the diagnostic window.

**Figure 5:**
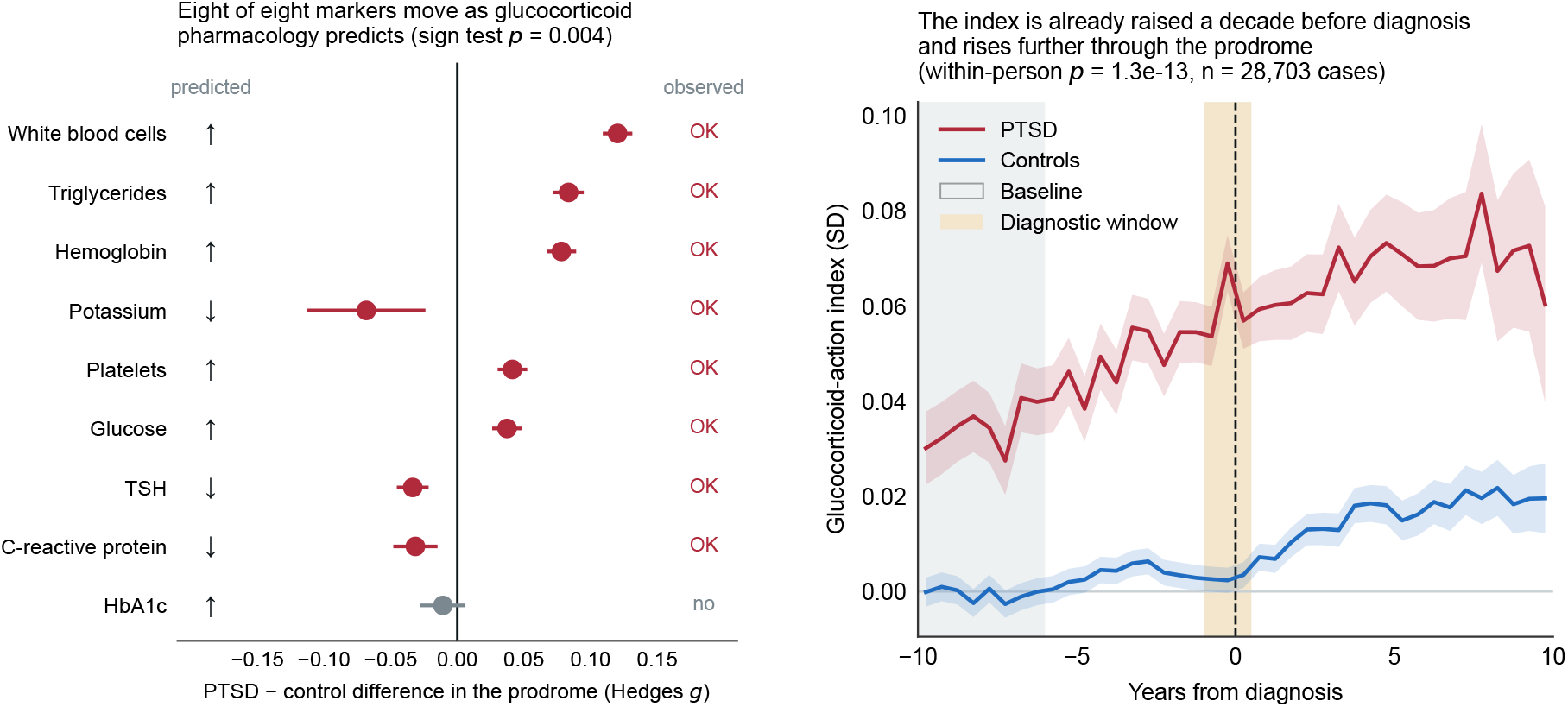
Peripheral glucocorticoid action in PTSD. (A) Hedges *g* with 95% CI for the PTSD − control difference in the prodrome, each marker’s direction assigned in advance from glucocorticoid pharmacology. Eight of the nine clear the 5% false-discovery threshold and all eight are concordant; HbA1c is the one marker that fails to move, its interval spanning zero. (B) The composite index across twenty years, person-level means in half-year bins with 95% CI bands; gray marks the baseline window and amber the diagnostic window, where differential testing surges and the level comparison should not be interpreted. Window means, confidence intervals and *n* are in Appendix Table S3.

C-reactive protein is the most informative marker in the set, because it separates glucocorrticoid action from general ill health: inflammation, obesity and illness all raise CRP, whereas glucocorticoid action lowers it (Michel et al., 2006; Vogel et al., 2023). CRP is indeed lower in PTSD in four of the five windows (*g* = −0.045 to −0.031, each passing the false-discovery rate). The one exception is the diagnostic window (*g* = +0.003, not passing it). That is the window in which differential testing surges, and in which people are drawn precisely because they are acutely unwell. A raised white-cell count together with a suppressed acute-phase response is the signature of glucocorticoid exposure, demargination neutrophilia without inflammation, rather than of an inflammatory state, and the two move that way together in a randomized placebo-controlled trial of dexamethasone (Barden et al., 2021). Markers on which glucocorticoid pharmacology makes no confident prediction (blood pressure, HDL, LDL and total cholesterol, transaminases, creatinine, albumin/creatinine) move, as a composite, slightly the other way in every window (−0.004 to −0.012 SD).

Standardizing each marker within sex against the control distribution, flipping it by its assigned sign and averaging over the markers a person carries in a half-year bin gives one glucocorticoid-action index per person per bin: 3.0 million person-bins, 418,000 of them from cases. The index is higher in PTSD in every window of the twenty-year record (Fig. 5B). Permuting arm labels across persons, which preserves the correlation between markers, gives *p* = 0.0005, the resolution limit of 2,000 permutations, in all five windows. Testing patterns change around diagnosis, so we also computed a within-person estimate that compares each person only with themselves: the change in each person’s index from their own baseline window, compared between arms. From baseline to prodrome, PTSD patients gain +0.0119 SD of glucocorticoid action relative to controls (*p* = 1.3 × 10^−13^; 28,703 cases have readings in both windows); the gain is +0.0214 SD by the diagnostic window and +0.0134 SD by the early window. Because each person serves as their own control, these estimates are insensitive to changes in who presents for testing. Excluding everyone with a tobacco record from both arms leaves the direction unchanged in the prodrome and early windows and shrinks the effects by about a quarter. Tobacco records are more common in cases (26.6% against 15.0%), so this quarter bounds the contribution of smoking from above.

The index is already displaced at baseline, a decade before diagnosis (+0.0375 SD, *g* = +0.105; Appendix Table S3), at 76% of the prodrome displacement. A difference that precedes diagnosis cannot be a consequence of PTSD; it is either an early glucocorticoid-specific shift or a prerexisting difference between the pools, such as adiposity or comorbidity. The non-directional control markers separate the two, because a gradient of general ill health would displace them at baseline alongside the directional markers. As a composite they move the other way at baseline (−0.0115 SD, *g* = −0.033), about a third the size of the directional offset, so the offset does not behave like a uniform gradient of morbidity. Individually, six of the nine non-directional markers clear the false-discovery threshold at baseline with signs that cancel, and the pattern (lower HDL and higher LDL, with white cells the largest directional effect) matches a smoking or metabolic signature, consistent with the excess of tobacco records among cases. The within-person gain, +0.0119 SD through the prodrome, is roughly a quarter of the cross-sectional contrast and is the estimate least sensitive to who presents for testing.

Cortisol is lower and peripheral glucocorticoid action is higher. Less hormone producing more effect in tissues outside the feedback loop requires a more sensitive receptor there, so the sensitization captured by *K*_GR_ is body-wide rather than confined to hypothalamus and pituitary, and a single *K*_GR_ accounts for the feedback and the peripheral findings together. The absence of a Cushingoid presentation follows from magnitude rather than site: a uniform *K*_GR_ raises occupancy by about a quarter of the rise in moderate Cushing’s syndrome (above), and the measured peripheral displacement is about a tenth of a standard deviation, a shift of population means rather than a clinical syndrome. The leukocyte data fit the same picture: circulating cells are peripheral tissue, and their 1.2–1.6-fold lower dexamethasone IC_50_ (Blalock et al., 2024; Rohleder et al., 2004; Yehuda et al., 2004b) is a sensitization of the expected sign, smaller than the feedback-arm ratio.

### 2.8 GR sensitization may precede the disorder

The model takes increased GR sensitivity as its input and is silent on its origin, but prospective data place it before the disorder: pre-deployment leukocyte GR number and pre-trauma GR-pathway markers predict later symptoms (van Zuiden et al., 2011, 2012), rats that later develop a PTSD-like phenotype already show stronger glucocorticoid feedback (Danan et al., 2021), and the small hippocampus of PTSD, the structural cost of sustained GR activation (Brown et al., 2015; Conrad, 2008; Kino, 2015; McEwen et al., 2016), is shared by unexposed co-twins (Gilbertson et al., 2002; Pitman et al., 2006). The glucocorticoid-action index displaced a decade before diagnosis points the same way. Trauma can also sensitize GR through *NR3C1* methylation (Labonté et al., 2014; Yehuda et al., 2014); the model separates the two origins, because only an acquired sensitization produces an ACTH deficit that appears after onset and resolves over about two months.

### 2.9 Predictions and limitations

The framework makes testable predictions: adrenal cortical volume ∼4% lower in PTSD, beyond the resolution of existing studies with 7–22 subjects per arm (Golier et al., 2014; Radant et al., 2009; Rasmusson et al., 2001); ACTH invariant across PTSD severity; the effect of cortisol synthesis blockers (*b*_3_) on basal cortisol lost over weeks as gland mass compensates, whereas CRH-directed agents act durably (Milo et al., 2025); GR antagonists durably raising basal cortisol while lowering GR occupancy, with MR signaling checked because MR is near saturation (de Kloet, 2022; Lembke et al., 2013; Otte et al., 2015); an induced change in GR sensitivity producing an ACTH deficit that resolves over about two months rather than persisting; and within-subject agreement between leukocyte and feedback GR sensitivity.

The peripheral screen has several limitations. Corticosteroid dispensing reproduces the marker signatures, and in the marker extract dispensing records are available for every case but for only 12% of controls, so no symmetric exclusion is possible; the cortisol cohort, a separate extract, has them for both groups. Where observed, exposure before the index date is strongly differential (38% of male and 49% of female cases against 15% and 23% of controls). In that subset, removing steroid-exposed people attenuates the cross-sectional index by a quarter to a third in the baseline and prodrome windows and leaves the within-person estimate unchanged; because the subset is not exchangeable with the full control pool, this is a bound, not a corrected estimate. The marker directions were not preregistered, and the sex-split and smoking analyses re-test the same cohort. Behavioral and metabolic sequelae of PTSD, including sleep, activity, diet and alcohol, could displace several of the markers without any receptor change, although the fall in CRP is difficult to obtain that way.

The model has limits. The TSST data are best fitted at a pre-stress drive fourfold above the operating point that population cortisol selects, a difference that basal and challenge cortisol measured in the same subjects would resolve. Some studies report disrupted ACTH, normal cortisol or intact challenge responses, so it may describe a subset of patients. The cortisol cohort is cross-sectional, from a single HMO, identifies PTSD from ICD codes, and cannot adjust for why a test was ordered or its exact draw time. The model lumps receptor number and affinity into *K*_GR_, does not represent binding-globulin capacity or 11*β*-HSD conversion, omits circadian and ultradian dynamics, and omits receptor-level compensation, which would attenuate the rise in GR occupancy, so the steady-state occupancy figures are upper bounds.

An endocrine axis whose glands adapt their functional mass hides most of its own parameters: integral feedback returns ACTH to its set point and absorbs changes in secretory capacity into gland size, leaving central drive and receptor sensitivity as the only changes basal cortisol can report. The low cortisol and normal ACTH of PTSD are what a sensitized receptor looks like in basal hormone levels once the glands have adapted, and the decisive measurement is therefore of glucocorticoid action rather than concentration. Read that way, routine clinical records point in one direction: cortisol in PTSD is not too low, but too high for the receptors that read it.

## 3 Methods

### 3.1 The HPA gland-mass model

The five-state model (Karin et al., 2020) is

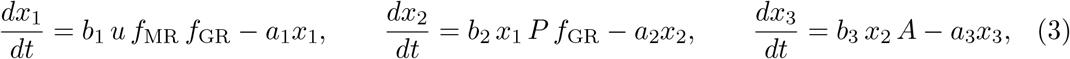

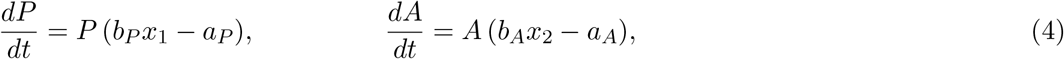

with *f*_GR_ = [1 + (*x*_3_*/K*_GR_)^3^]^−1^ and *f*_MR_ = 1*/x*_3_, the latter a local approximation valid near baseline cortisol. The Hill coefficient *n* = 3 is taken from the published calibration (Karin et al., 2020); it is a lumped exponent standing for receptor dimerization and multi-site transcriptional activation. The PTSD occupancy for *n* = 2 and *n* = 4 is given in the Results, and the cortisol reduction for each in Fig. 4C. Rate constants follow the published calibration (Karin et al., 2020) (*a*_1_ = 0.17, *a*_2_ = 0.035 min^−1^; corticotroph and adrenal turnover 0.099 and 0.049 day^−1^), except cortisol clearance, set to *a*_3_ = 0.06 min^−1^ to reflect the shorter free-cortisol disposition kinetics reported by Dorin et al. (2022). Secretion equals degradation for each species, normalizing every baseline steady state to 1. Only *K*_GR_ differs between PTSD and control simulations (Appendix Table S1).

Eliminating *x*_1_ and *x*_2_ from Eqs. (3) with *P* = *A* = 1, and noting that *f*_GR_ enters both the CRH and ACTH conditions, gives *x*_3_ = *Cf*_GR_ with 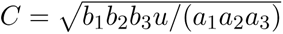, i.e. 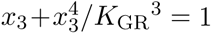 for *C* = 1. Equation (2) is the corresponding relation for the full model with adaptive masses, where *C* is replaced by *b*_1_*b_P_ u/*(*a*_1_*a_P_*); the two coincide under the normalization used here, and this is the curve swept in Fig. 4C,D.

### 3.2 Numerical simulations

Equations were integrated with `scipy.integrate.odeint` (LSODA, default tolerances) from the pre-stimulus steady state, with `hmax`= 0.01 for the minute-scale challenge simulations and `hmax`= 5 for the 180-day onset transient; steady states were found with `scipy.optimize.fsolve`. Over the range considered the steady-state condition has a unique root in (0, 1), so the model is monostable.

The TSST was simulated as a 20-minute tenfold increase in *u*, from a pre-stress value of 4 to 40, initialized at the steady state for *u* = 4 and plotted relative to that baseline; the dimensionless steady-state scans and the Dex–CRH simulation use *u* = 1.

The pre-stress level was chosen by SEM-weighted fit to the measured time courses, over pre-stress levels from 0.5 to 15 and pulses of three- to twentyfold. *u* = 4 and *u* = 3 fit best (*χ*^2^ per point 0.77), and *u* = 4 places more points within their SEM (10 of 15). With *u* held at 1, the best fit over pulse sizes and durations gives *χ*^2^ per point 3.3, because the pre-stress difference is then fixed at the 4.4% steady-state value. In the published means of von Majewski et al. (2023), pre-stress cortisol in the PTSD arm is about 30% below control, its peak about 50% below, and its rise above its own baseline about 30% of the control rise; the *u* = 4 axis gives 28%, 37% and 55% for these three quantities, and the *u* = 1 axis 4%, 16% and 78%. Over the pre-stress levels and pulse sizes tested, the peak reduction stays below ∼40% and the rise ratio above ∼55%.

The two Dex–CRH input magnitudes, shared by both arms, were set by SEM-weighted least squares against the measured ACTH with *K*_GR_ fixed at 2.5/5. Dexamethasone enters as a cortisol-equivalent GR agonist *D* = 7 added to *x*_3_ inside the GR Hill term, 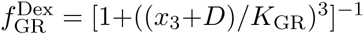, and is held constant from the start of the run; because the agonist is not cleared at the cortisol rate the suppressed state is reached within a few hours, so it is applied ∼7 h ahead of the CRH stimulus. Dexamethasone is excluded from the CRH equation, since it crosses the blood–brain barrier minimally. Exogenous CRH is a 20-minute pulse of magnitude *u*_CRH_ = 0.35 added to *dx*_1_*/dt*, beginning at the plotted *t* = 0. The CRH-arm rate constants are rescaled by 0.1 (*b*_1_ = *a*_1_ = 0.017 min^−1^) to set the CRH timescale of the challenge. Only *K*_GR_ differs between the two arms.

#### Dexamethasone suppression

To compare the calibration with dexamethasone suppression data, the Dex term 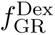 defined above, with a strength *D* set separately from the Dex–CRH value, was switched on at 23:00 in an axis at its baseline steady state and cortisol was read 9 h later, at 08:00; gland masses were left free and change by less than 10% over this interval. The Dex strength *D* was set so that suppression in the control arm (*K*_GR_ = 5) equals the observed value (72–78%; Newport et al., 2004; Yehuda et al., 2002); the quantity tested is the PTSD arm (*K*_GR_ = 2.5) at the same *D*. The model then predicts 88–91% suppression in PTSD, a 2.3-fold lower residual cortisol, against 2.4-fold in PTSD without comorbid depression and 1.5-fold in all PTSD subjects (Yehuda et al., 2002), and 3.5-fold in abuse survivors with PTSD (Newport et al., 2004).

Both challenge panels plot the model on the same axes as the measurements (Fig. 3). The model is normalized, so a display factor is needed to carry it into the measured units; one factor per panel is fitted by least squares over every point of both arms at once, which sets only the absolute scale. For the Dex–CRH panel this factor is ≈ 30 pg/mL per model unit. For the TSST panel it is 0.74 nmol/L per model unit. The measurements were digitized from the published panels of von Majewski et al. (2023) and Ströhle et al. (2008) by calibrating each axis from its tick-mark pixels and locating each marker by template-matching the marker glyph, with error-bar caps read from their own pixel runs; both papers draw one-sided bars, so the plotted intervals are the published SEM. Digitization error is below one tenth of a SEM.

### 3.3 Observability and site-resolved analysis

The observability statement was obtained symbolically from (Eqs. 3)–(4) and confirmed numerircally by perturbing each parameter by ±20% and re-solving the fixed point. The site-resolved analysis replaces the single *K*_GR_ with *K*_GR_^hyp^ in *dx*_1_*/dt* and *K*_GR_^pit^ in *dx*_2_*/dt*. The drive-versus-sensitivity comparison solves for the reduction in *u* and the reduction in *K*_GR_ that each produce the same cortisol fall, and evaluates GR occupancy at both solutions. The site-resolved split is a limiting case: the peripheral marker screen selects body-wide sensitization, so *K*_GR_^hyp^ and *K*_GR_^pit^ are assumed equal in all other simulations, and the dexamethasone comparison made for the pituitary arm is taken to hold for the hypothalamic arm on that basis.

### 3.4 Morning cortisol analysis

Morning serum cortisol tests (06:00–10:00, ages 18–70) were extracted from Clalit electronic medical records as pseudonymized per-test records. PTSD cases carried ICD-9 309.81 or ICD-10 F43.1, and only their tests taken after the first PTSD diagnosis were used; control tests were taken at any time. Individuals with diagnoses affecting HPA function (ICD-9 rubrics 253, 255 and 296, with all subcodes) were excluded from both groups. In both groups, a test was excluded if any of 14 corticosteroids (any formulation) had been dispensed in the 90 days before it; individuals with other, unaffected tests were retained. Tests were aggregated into 3-year age bins; group means were compared by direct age standardization to the combined unique-individual age distribution, with significance from a two-sample *z*-test on standardized means and standard errors propagated from per-bin values. Hedges’ *g* confidence intervals were obtained as *g* ± 1.96 |*g*|*/z*. Random-effects meta-analysis of per-bin differences gives men *p* = 0.031, women *p* = 0.020. Test-per-person ratios are 1.2–1.3, and an analysis with one test per individual gives men *p* = 0.025, women *p* = 0.002.

### 3.5 Peripheral laboratory-marker screen

Nine glucocorticoid-responsive analytes, together with a further set carrying no confident directional prediction, were extracted for 40,240 PTSD cases and 311,362 controls from the same records: 26.1 million readings of the nine directional analytes within ±10 y of the index date, and 52.0 million across all eighteen. Cases carried the same ICD-9 309.81 or ICD-10 F43.1 codes as the cortisol cohort, with a first recorded diagnosis between ages 20 and 80, and their index date is that first diagnosis. Controls are members with no PTSD code at any time and at least one recorded visit, matched to cases on sex, year of birth and primary-care clinic: within each clinic-by-birth-year stratum, controls were drawn at random without replacement from the full member pool, up to ten per case; clinic was required to be recorded. Each control takes as its index the diagnosis age of a case in its stratum. Because birth year is matched, a control’s index falls in approximately the same calendar year as its case’s diagnosis, and readings from both arms are placed on the same years-from-index axis (Fig. 5B). The screen retains 7.7 controls per case. Marker comparisons are not adjusted for age; cases are 0.7–0.9 y older, and re-standardizing within five-year age bands changes the composite index by less than 5% in every window. Directions were assigned from the pharmacological literature before any group comparison. Analytes for which the literature supports no single direction were screened without a directional assignment and are excluded from the sign test. Each assay is filtered to a physiological window; triglycerides and TSH are log-transformed before standardization. Each reading is z-scored within marker and sex against the control distribution and multiplied by its assigned sign, then averaged within person within half-year bin and across the markers that person carries in that bin; bins with fewer than four markers are dropped. Window statistics take one value per person per window and compare arms with Welch’s *t*; *g* is Hedges’ *g*; permutation *p* shuffles arm labels across persons within sex, 2,000 draws. The false-discovery correction is applied within window, across the nine markers; sign-test *p* values are one-sided. The within-person estimator subtracts each person’s own −10 to −6 y index. Window boundaries follow a broken-stick breakpoint analysis of the same cohort, which places the baseline ahead of every observed turn and sets aside −1 to +0.5 y, where differential testing surges.

## Data availability

The computer code produced in this study is available in the following database:

- Model code (the equations, the simulations, the observability analysis, the digitized challenge-test measurements and the regression tests): Zenodo, doi:10.5281/zenodo.22978219 (Danan et al., 2026), the archived release v1.0.0 of https://github.com/AlonLabWIS/hpa-dysregulation-ptsd.

The code reproduces Figs. 2 and 3, the model panels of Fig. 4C,D and every model number quoted in the text from a clean checkout with no external data, and its regression tests assert those numbers. The cohort results (Fig. 4A,B and Fig. 5) are computed from Clalit Health Services electronic health records, which cannot be redistributed under the terms of the data-governance agreement; that pipeline therefore cannot be run outside Clalit, and access requests are made to Clalit.

## Ethics approval

Use of the de-identified Clalit electronic-medical-record data was approved by the Clalit Helsinki (institutional review board) Committee (approval RMC-1059-21), which covers both the cortisol and the laboratory-marker cohorts and waived the requirement for informed consent for this retrospective analysis of de-identified data. The human stress-test and Dex–CRH data shown for comparison (Ströhle et al., 2008; von Majewski et al., 2023) are reproduced from previously published studies.

## Author contributions

D.D. and Y.G. contributed equally. **Dor Danan:** Conceptualization, Methodology, Software, Formal analysis, Writing — original draft, Writing — review & editing. **Yaniv Grosskopf:** Methodology, Software, Formal analysis, Writing — original draft, Writing — review & editing. **Yoav Hayut:** Methodology, Software, Writing — review & editing. **Yoel Toledano:** Resources, Data curation, Writing — review & editing. **Keren Doenyas-Barak:** Resources, Data curation, Writing — review & editing. **Avi Mayo:** Conceptualization, Methodology, Supervision, Writing — review & editing. **Uri Alon:** Conceptualization, Resources, Supervision, Funding acquisition, Writing — review & editing.

## Acknowledgements

We thank all members of our lab, Amos Tanay, Neta Mendelsohn and Rami Jaschek for discussions, and Gabi Barabash and Ran Balicer for the Clalit–Weizmann collaboration.

## Appendix

### Appendix Table S1. Model variables and parameters

**Appendix Table S1.**
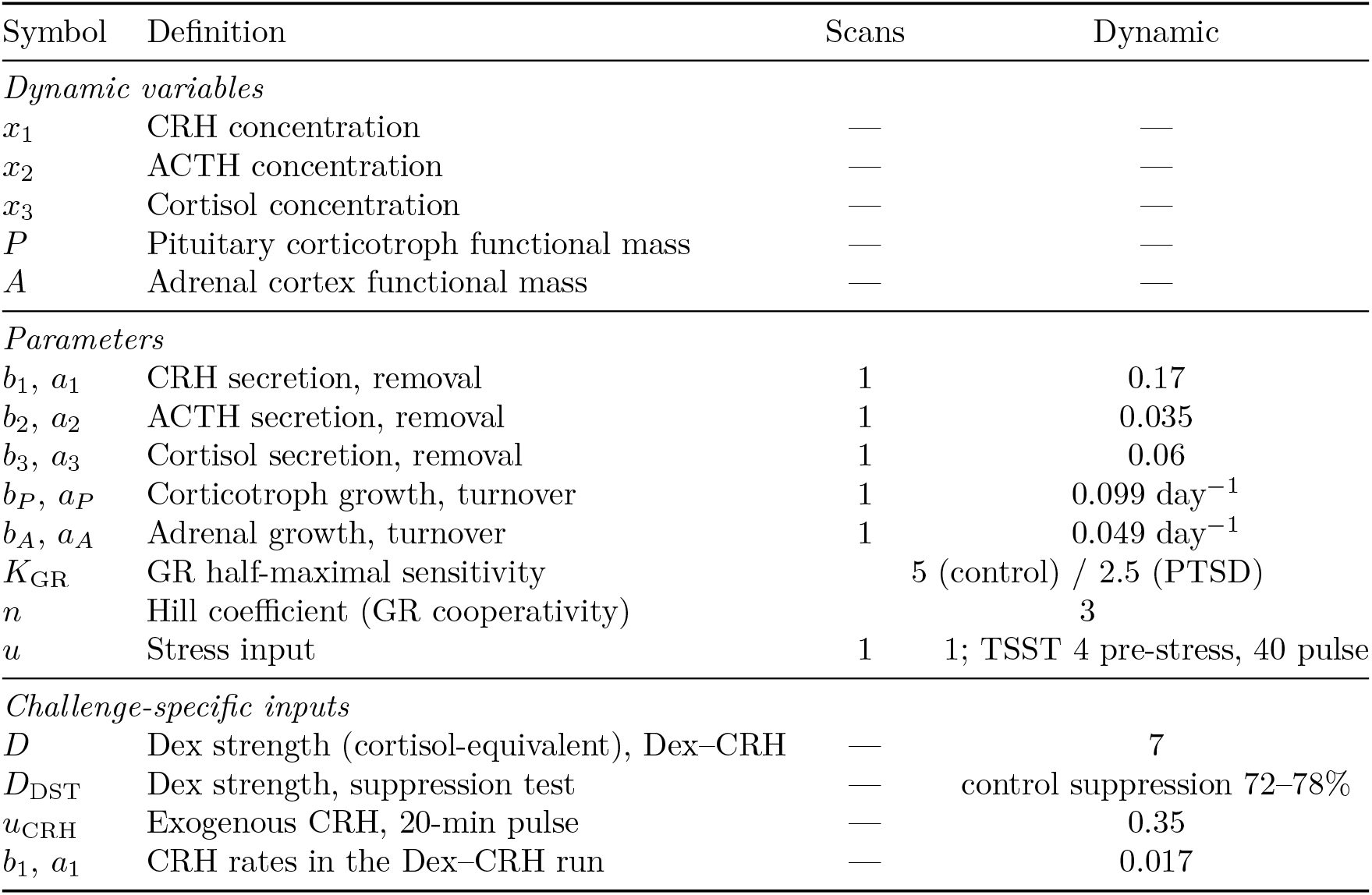
Variables and parameters of the HPA gland-mass model. Hormone rates are in min^−1^ and gland-mass turnover rates in day^−1^ (converted to min^−1^ in the simulations). Because secretion equals degradation for every species, each baseline steady state is normalized to 1, so the steady-state scans use unit rates while the dynamic simulations use the dimensional values. *K*_GR_ is the only quantity that differs between PTSD and control. The challenge-specific inputs apply to the named simulation only.

### Appendix Table S2. Morning serum cortisol by sex and PTSD status

**Appendix Table S2.**
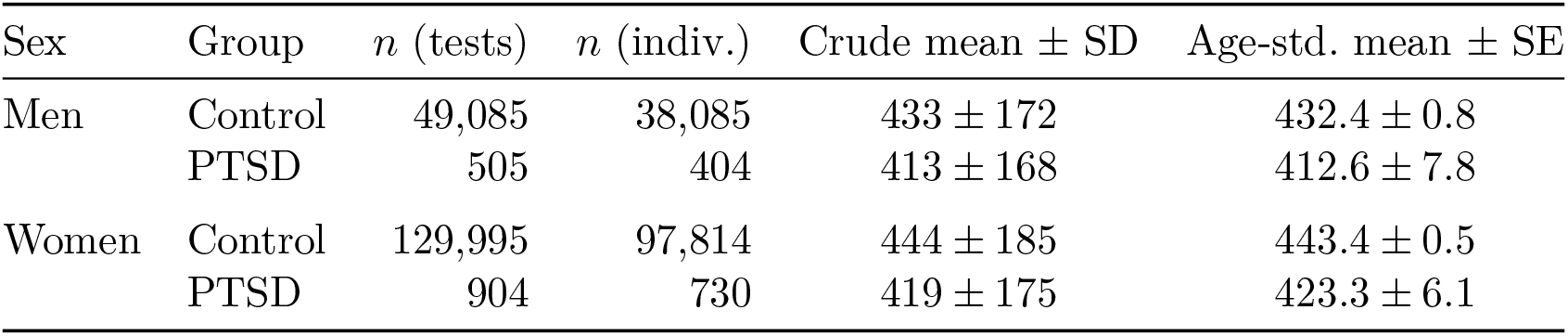
Crude and age-standardized morning serum cortisol (nmol/L) by sex and PTSD status, adults aged 18–70. Standardization uses the combined PTSD-plus-control unique-individual age distribution as the common reference, so the standardized means differ only by the PTSD effect and not by age structure. Standardized contrasts: men Δ = *−*19.8 nmol/L (*−*4.6%), *g* = *−*0.11 (95% CI *−*0.20 to *−*0.03), *p* = 0.012; women Δ = *−*20.1 nmol/L (*−*4.5%), *g* = *−*0.11 (*−*0.17 to *−*0.04), *p* = 0.001. Unique individuals are counted once each across age bins.

| Sex | Group | $n$ (tests) | $n$ (indiv.) | Crude mean ± SD | Age-std. mean ± SE |
| --- | --- | --- | --- | --- | --- |
| Men | Control | 49,085 | 38,085 | $433 \pm 172$ | $432.4 \pm 0.8$ |
| | PTSD | 505 | 404 | $413 \pm 168$ | $412.6 \pm 7.8$ |
| Women | Control | 129,995 | 97,814 | $444 \pm 185$ | $443.4 \pm 0.5$ |
| | PTSD | 904 | 730 | $419 \pm 175$ | $423.3 \pm 6.1$ |

### Appendix Table S3. Glucocorticoid-action index by window

**Appendix Table S3.**
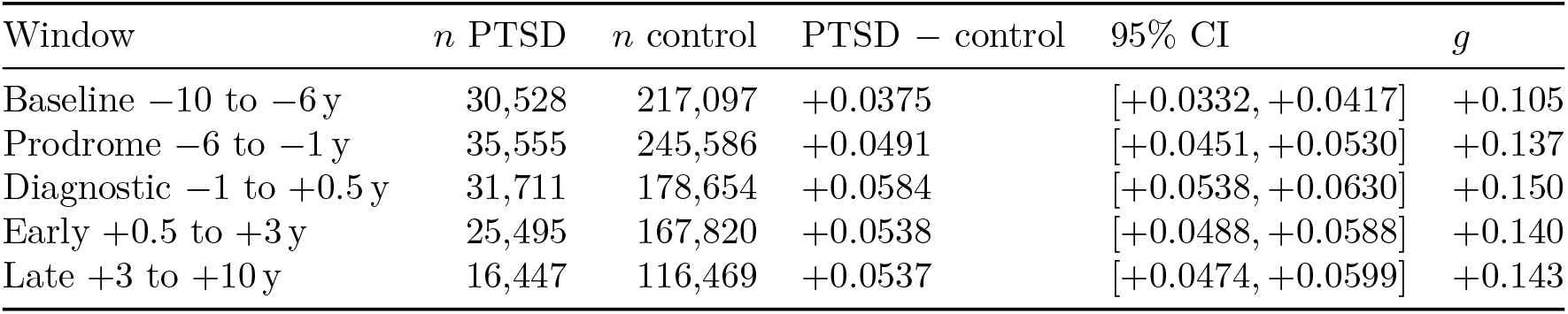
Glucocorticoid-action index, in SD units of the control distribution. Person-level: each person contributes one value per window. *p* from 2,000 permutations of arm labels within sex; 0.0005 is the resolution limit. PTSD and control columns give the number of persons.

## References

M. Algamal et al. Chronic Hippocampal Abnormalities and Blunted HPA Axis in an Animal Model of Repeated Unpredictable Stress. Front. Behav. Neurosci., 12:150, 2018. doi: 10.3389/fnbeh.2018.00150.

S. L. Asa et al. Pituitary corticotroph hyperplasia in rats implanted with a medullary thyroid carcinoma cell line transfected with a corticotropin-releasing hormone complementary deoxyribonucleic acid expression vector. Endocrinology, 131:715–720, 1992. doi: 10.1210/endo.131.2.1322279.

M. Atmaca, O. Ozer, S. Korkmaz, I. Taskent, and H. Yildirim. Evidence for the changes of pituitary volumes in patients with post-traumatic stress disorder. Psychiatry Res. Neuroimaging, 260:49–52, 2017. doi: 10.1016/j.pscychresns.2016.12.004.

R. D. Balicer and A. Afek. Digital health nation: Israel’s global big data innovation hub. Lancet, 389:2451–2453, 2017. doi: 10.1016/s0140-6736(17)30876-0.

A. Bar et al. Pregnancy and postpartum dynamics revealed by millions of lab tests. Sci. Adv., 11:eadr7922, 2025. doi: 10.1126/sciadv.adr7922.

A. Barden, M. Phillips, S. Shinde, T. Corcoran, and T. A. Mori. The effects of perioperative dexamethasone on eicosanoids and mediators of inflammation resolution: a sub-study of the PADDAG trial. Prostaglandins Leukot. Essent. Fatty Acids, 173:102334, 2021. doi: 10.1016/j.plefa.2021.102334.

M. E. Bauer, L. K. Price, M. P. MacEachern, M. Housey, E. S. Langen, and S. T. Bauer. Maternal leukocytosis after antenatal corticosteroid administration: a systematic review and meta-analysis. J. Obstet. Gynaecol., 38(2):210–216, 2018. doi: 10.1080/01443615.2017.1342614.

E. M. Berntsen, M. D. Haukedal, and A. K. Håberg. Normative data for pituitary size and volume in the general population between 50 and 66 years. Pituitary, 24:737–745, 2021. doi: 10.1007/s11102-021-01150-7.

F. Beuschlein et al. European Society of Endocrinology and Endocrine Society Joint Clinical Guideline: Diagnosis and Therapy of Glucocorticoid-induced Adrenal Insufficiency. J. Clin. Endocrinol. Metab., 109:1657–1683, 2024. doi: 10.1210/clinem/dgae250.

Z. N. Blalock, G. W. Y. Wu, D. Lindqvist, C. Trumpff, J. D. Flory, J. Lin, V. I. Reus, R. Rampersaud, R. Hammamieh, A. Gautam, F. J. Doyle, C. R. Marmar, M. Jett, R. Yehuda, O. M. Wolkowitz, and S. H. Mellon. Circulating cell-free mitochondrial DNA levels and glucocorticoid sensitivity in a cohort of male veterans with and without combat-related PTSD. Transl. Psychiatry, 14:22, 2024. doi: 10.1038/s41398-023-02721-x.

E. S. Brown et al. Hippocampal Volume in Healthy Controls Given 3-Day Stress Doses of Hydrocortisone. Neuropsychopharmacology, 40:1216–1221, 2015. doi: 10.1038/npp.2014.307.

T. O. Bruhn, R. E. Sutton, C. L. Rivier, and W. W. Vale. Corticotropin-releasing factor regulates proopiomelanocortin messenger ribonucleic acid levels in vivo. Neuroendocrinology, 39:170–175, 1984. doi: 10.1159/000123974.

R. R. Cavalieri. The effects of nonthyroid disease and drugs on thyroid function tests. Med. Clin. North Am., 75(1):27–39, 1991. doi: 10.1016/s0025-7125(16)30470-9.

J.-H. Cho and S. Suh. Glucocorticoid-induced hyperglycemia: a neglected problem. Endocrinol. Metab. (Seoul), 39(2):222–238, 2024. doi: 10.3803/EnM.2024.1951.

N. M. Cohen et al. Personalized lab test models to quantify disease potentials in healthy individuals. Nat. Med., 27:1582–1591, 2021. doi: 10.1038/s41591-021-01468-6.

C. D. Conrad. Chronic stress-induced hippocampal vulnerability: the glucocorticoid vulnerability hypothesis. Rev. Neurosci., 19:395–411, 2008. doi: 10.1515/revneuro.2008.19.6.395.

O. Cooper, V. Bonert, F. Moser, J. Mirocha, and S. Melmed. Altered Pituitary Gland Structure and Function in Posttraumatic Stress Disorder. J. Endocr. Soc., 1:577–587, 2017. doi: 10.1210/js.2017-00069.

A. D. Crockard, M. T. Boylan, A. G. Droogan, S. A. McMillan, and S. A. Hawkins. Methylprednisolone-induced neutrophil leukocytosis: down-modulation of neutrophil L-selectin and Mac-1 expression and induction of granulocyte-colony stimulating factor. Int. J. Clin. Lab. Res., 28(2):110–115, 1998. doi: 10.1007/s005990050029.

D. Danan, D. Todder, J. Zohar, and H. Cohen. Is PTSD-Phenotype Associated with HPA-Axis Sensitivity? Feedback Inhibition and Other Modulating Factors of Glucocorticoid Signaling Dynamics. Int. J. Mol. Sci., 22(11):6050, 2021. doi: 10.3390/ijms22116050.

D. Danan, Y. Grosskopf, Y. Hayut, Y. Toledano, K. Doenyas-Barak, A. Mayo, and U. Alon. Model code for “Gland-Mass Adaptation Explains the Cortisol Paradox in PTSD”. Zenodo, version v1.0.0, 2026. URL 10.5281/zenodo.22978219.

C. S. de Kloet, E. Vermetten, A. Bikker, E. Meulman, E. Geuze, A. Kavelaars, H. G. M. Westenberg, and C. J. Heijnen. Leukocyte glucocorticoid receptor expression and immunoregulation in veterans with and without post-traumatic stress disorder. Mol. Psychiatry, 12(5):443–453, 2007. doi: 10.1038/sj.mp.4001934.

C. S. de Kloet et al. Assessment of HPA-axis function in posttraumatic stress disorder: pharmacological and non-pharmacological challenge tests, a review. J. Psychiatr. Res., 40:550–567, 2006. doi: 10.1016/j.jpsychires.2005.08.002.

E. R. de Kloet. Brain mineralocorticoid and glucocorticoid receptor balance in neuroendocrine regulation and stress-related psychiatric etiopathologies. Curr. Opin. Endocr. Metab. Res., 24: 100352, 2022. doi: 10.1016/j.coemr.2022.100352.

R. I. Dorin, F. K. Urban, I. Perogamvros, and C. R. Qualls. Four-compartment diffusion model of cortisol disposition: Comparison to three alternative models in current clinical use. J. Endocr. Soc., 7(1):bvac173, 2022. doi: 10.1210/jendso/bvac173.

M. W. Gilbertson et al. Smaller hippocampal volume predicts pathologic vulnerability to psychological trauma. Nat. Neurosci., 5:1242–1247, 2002. doi: 10.1038/nn958.

Julia A. Golier, Kimberly Caramanica, and Rachel Yehuda. Neuroendocrine response to CRF stimulation in veterans with and without PTSD in consideration of war zone era. Psychoneuroendocrinology, 37(3):350–357, 2012. doi: 10.1016/j.psyneuen.2011.07.004.

Julia A. Golier, Kimberly Caramanica, Iouri Makotkine, Leo Sher, and Rachel Yehuda. Cortisol response to cosyntropin administration in military veterans with or without posttraumatic stress disorder. Psychoneuroendocrinology, 40:151–158, 2014. doi: 10.1016/j.psyneuen.2013.10.020.

J. Herndon, R. J. Kaur, M. Romportl, E. Smith, A. Koenigs, B. Partlow, L. Arteaga, and I. Bancos. The effect of curative treatment on hyperglycemia in patients with Cushing syndrome. J. Endocr. Soc., 6(1):bvab169, 2021. doi: 10.1210/jendso/bvab169.

A. M. Isidori, M. Minnetti, E. Sbardella, C. Graziadio, and A. B. Grossman. Mechanisms in endocrinology: The spectrum of haemostatic abnormalities in glucocorticoid excess and defect. Eur. J. Endocrinol., 173(3):R101–R113, 2015. doi: 10.1530/EJE-15-0308.

E. D. Kanter et al. Glucocorticoid feedback sensitivity and adrenocortical responsiveness in posttraumatic stress disorder. Biol. Psychiatry, 50:238–245, 2001. doi: 10.1016/s0006-3223(01)01158-1.

O. Karin, A. Swisa, B. Glaser, Y. Dor, and U. Alon. Dynamical compensation in physiological circuits. Mol. Syst. Biol., 12(11):886, 2016. doi: 10.15252/msb.20167216.

O. Karin, M. Raz, and U. Alon. An opponent process for alcohol addiction based on changes in endocrine gland mass. iScience, 24:102127, 2021. doi: 10.1016/j.isci.2021.102127.

O. Karin et al. A new model for the HPA axis explains dysregulation of stress hormones on the timescale of weeks. Mol. Syst. Biol., 16:e9510, 2020. doi: 10.15252/msb.20209510.

L. U. Kim, M. R. D’Orsogna, and T. Chou. Onset, timing, and exposure therapy of stress disorders: mechanistic insight from a mathematical model of oscillating neuroendocrine dynamics. Biol. Direct, 11:13, 2016. doi: 10.1186/s13062-016-0117-6.

L. U. Kim, M. R. D’Orsogna, and T. Chou. Perturbing the Hypothalamic–Pituitary–Adrenal Axis: A Mathematical Model for Interpreting PTSD Assessment Tests. *Comput*. Psychiatry, 2:28, 2018. doi: 10.1162/CPSY_a_00013.

T. Kino. Stress, glucocorticoid hormones, and hippocampal neural progenitor cells: implications to mood disorders. Front. Physiol., 6:230, 2015. doi: 10.3389/fphys.2015.00230.

B. Labonté, N. Azoulay, V. Yerko, G. Turecki, and A. Brunet. Epigenetic modulation of glucocorticoid receptors in posttraumatic stress disorder. Transl. Psychiatry, 4:e368, 2014. doi: 10.1038/tp.2014.3.

S. Lawrence and R. H. Scofield. Post traumatic stress disorder associated hypothalamic-pituitary-adrenal axis dysregulation and physical illness. Brain Behav. Immun. - Health, 41:100849, 2024. doi: 10.1016/j.bbih.2024.100849.

A. Lembke et al. The mineralocorticoid receptor agonist, fludrocortisone, differentially inhibits pituitary-adrenal activity in humans with psychotic major depression. Psychoneuroendocrinology, 38:115–121, 2013. doi: 10.1016/j.psyneuen.2012.05.006.

D. G. Levitt. Pharmacokinetics/pharmacodynamics of glucocorticoids: modeling the glucocorticoid receptor dynamics and dose/response of commonly prescribed glucocorticoids. ADMET DMPK, 12:971–989, 2024. doi: 10.5599/admet.2414.

C. F. P. Lotfi and P. O. R. De Mendonca. Comparative Effect of ACTH and Related Peptides on Proliferation and Growth of Rat Adrenal Gland. Front. Endocrinol., 7:39, 2016. doi: 10.3389/fendo.2016.00039.

E. B. Manukhina et al. Intermittent hypoxia improves behavioral and adrenal gland dysfunction induced by posttraumatic stress disorder in rats. J. Appl. Physiol., 125:931–937, 2018. doi: 10.1152/japplphysiol.01123.2017.

G. Matić, D. V. Milutinović, J. Nestorov, I. Elaković, S. M. Jovanović, T. Perišić, J. Dunđerski, S. Damjanović, G. Knežević, Ž. Špirić, E. Vermetten, and D. Savić. Lymphocyte glucocorticoid receptor expression level and hormone-binding properties differ between war trauma-exposed men with and without PTSD. Prog. Neuropsychopharmacol. Biol. Psychiatry, 43:238–245, 2013. doi: 10.1016/j.pnpbp.2013.01.005.

G. Matić, D. V. Milutinović, I. Elaković, J. Nestorov, and D. Savić. Level of Expression and Functional Properties of Lymphocyte Corticosteroid Receptors as Biological Correlates of PTSD, Trauma-Exposure or Resilience to PTSD. In C. R. Martin, V. R. Preedy, and V. B. Patel, editors, Comprehensive Guide to Post-Traumatic Stress Disorder, pages 1–16. Springer International Publishing, Cham, 2015. doi: 10.1007/978-3-319-08613-2_3-1.

B. S. McEwen, C. Nasca, and J. D. Gray. Stress Effects on Neuronal Structure: Hippocampus, Amygdala, and Prefrontal Cortex. Neuropsychopharmacology, 41:3–23, 2016. doi: 10.1038/npp.2015.171.

A. C. McFarlane, C. A. Barton, R. Yehuda, and G. Wittert. Cortisol response to acute trauma and risk of posttraumatic stress disorder. Psychoneuroendocrinology, 36:720–727, 2011. doi: 10.1016/j.psyneuen.2010.10.007.

M.-L. Meewisse, J. B. Reitsma, G.-J. de Vries, B. P. R. Gersons, and M. Olff. Cortisol and posttraumatic stress disorder in adults: systematic review and meta-analysis. Br. J. Psychiatry, 191:387–392, 2007. doi: 10.1192/bjp.bp.106.024877.

S. Melmed et al. Clinical Biology of the Pituitary Adenoma. Endocr. Rev., 43:1003–1037, 2022. doi: 10.1210/endrev/bnac010.

S. Metz, M. Duesenberg, J. Hellmann-Regen, O. T. Wolf, S. Roepke, C. Otte, and K. Wingenfeld. Blunted salivary cortisol response to psychosocial stress in women with posttraumatic stress disorder. J. Psychiatr. Res., 130:112–119, 2020. doi: 10.1016/j.jpsychires.2020.07.014.

O. Michel, M. Dentener, D. Cataldo, B. Cantinieaux, F. Vertongen, C. Delvaux, and R. D. Murdoch. Evaluation of oral corticosteroids and phosphodiesterase-4 inhibitor on the acute inflammation induced by inhaled lipopolysaccharide in human. Pulm. Pharmacol. Ther., 20 (6):676–683, 2006. doi: 10.1016/j.pupt.2006.08.002.

T. Milo et al. Hormone circuit explains why most HPA drugs fail for mood disorders and predicts the few that work. Mol. Syst. Biol., 21:254–273, 2025. doi: 10.1038/s44320-024-00083-0.

D. J. Newport, C. Heim, R. Bonsall, A. H. Miller, and C. B. Nemeroff. Pituitary-adrenal responses to standard and low-dose dexamethasone suppression tests in adult survivors of child abuse. Biol. Psychiatry, 55(1):10–20, 2004. doi: 10.1016/s0006-3223(03)00692-9.

L. Nolan and A. Levy. Anterior pituitary trophic responses to dexamethasone withdrawal and repeated dexamethasone exposures. J. Endocrinol., 169:263–270, 2001. doi: 10.1677/joe.0.1690263.

C. Otte et al. Mineralocorticoid receptor stimulation improves cognitive function and decreases cortisol secretion in depressed patients and healthy individuals. Neuropsychopharmacology, 40: 386–393, 2015. doi: 10.1038/npp.2014.181.

X. Pan, Z. Wang, X. Wu, S. W. Wen, and A. Liu. Salivary cortisol in post-traumatic stress disorder: a systematic review and meta-analysis. BMC Psychiatry, 18:324, 2018. doi: 10.1186/s12888-018-1910-9.

X. Pan et al. The 24-hour urinary cortisol in post-traumatic stress disorder: A meta-analysis. PLoS One, 15:e0227560, 2020. doi: 10.1371/journal.pone.0227560.

R. K. Pitman et al. Clarifying the origin of biological abnormalities in PTSD through the study of identical twins discordant for combat exposure. Ann. N. Y. Acad. Sci., 1071:242–254, 2006. doi: 10.1196/annals.1364.019.

M. Quinkler and P. M. Stewart. Hypertension and the cortisol-cortisone shuttle. J. Clin. Endocrinol. Metab., 88(6):2384–2392, 2003. doi: 10.1210/jc.2003-030138.

Allen D. Radant, Dorcas J. Dobie, Elaine R. Peskind, M. Michele Murburg, Eric C. Petrie, Evan D. Kanter, Murray A. Raskind, and Charles W. Wilkinson. Adrenocortical responsiveness to infusions of physiological doses of ACTH is not altered in posttraumatic stress disorder. Frontiers in Behavioral Neuroscience, 3:40, 2009. doi: 10.3389/neuro.08.040.2009.

A. M. Rasmusson et al. Increased pituitary and adrenal reactivity in premenopausal women with posttraumatic stress disorder. Biol. Psychiatry, 50:965–977, 2001. doi: 10.1016/s0006-3223(01)01264-1.

N. Rohleder, L. Joksimovic, J. M. Wolf, and C. Kirschbaum. Hypocortisolism and increased glucocorticoid sensitivity of pro-Inflammatory cytokine production in Bosnian war refugees with posttraumatic stress disorder. Biol. Psychiatry, 55:745–751, 2004. doi: 10.1016/j.biopsych.2003.11.018.

N. Rohleder, J. M. Wolf, and O. T. Wolf. Glucocorticoid sensitivity of cognitive and inflammatory processes in depression and posttraumatic stress disorder. Neurosci. Biobehav. Rev., 35:104–114, 2010. doi: 10.1016/j.neubiorev.2009.12.003.

A. Roussel-Gervais et al. The Cables1 Gene in Glucocorticoid Regulation of Pituitary Corticotrope Growth and Cushing Disease. J. Clin. Endocrinol. Metab., 101:513–522, 2016. doi: 10.1210/jc.2015-3324.

P. R. Somvanshi et al. Role of enhanced glucocorticoid receptor sensitivity in inflammation in PTSD: insights from computational model for circadian-neuroendocrine-immune interactions. Am. J. Physiol. Endocrinol. Metab., 319:E48–E66, 2020. doi: 10.1152/ajpendo.00398.2019.

K. Sriram, M. Rodriguez-Fernandez, and F. J. Doyle. Modeling cortisol dynamics in the neuro-endocrine axis distinguishes normal, depression, and post-traumatic stress disorder (PTSD) in humans. PLoS Comput. Biol., 8:e1002379, 2012. doi: 10.1371/journal.pcbi.1002379.

A. Ströhle, M. Scheel, S. Modell, and F. Holsboer. Blunted ACTH response to dexamethasone suppression-CRH stimulation in posttraumatic stress disorder. J. Psychiatr. Res., 42:1185–1188, 2008. doi: 10.1016/j.jpsychires.2008.01.015.

H. G. Swann. The Pituitary-Adrenocortical Relationship. Physiol. Rev., 20:493–521, 1940. doi: 10.1152/physrev.1940.20.4.493.

M. R. Taskinen, E. A. Nikkilä, R. Pelkonen, and T. Sane. Plasma lipoproteins, lipolytic enzymes, and very low density lipoprotein triglyceride turnover in Cushing’s syndrome. J. Clin. Endocrinol. Metab., 57(3):619–626, 1983. doi: 10.1210/jcem-57-3-619.

A. Tendler et al. Hormone seasonality in medical records suggests circannual endocrine circuits. Proc. Natl. Acad. Sci. U. S. A., 118:e2003926118, 2021. doi: 10.1073/pnas.2003926118.

L. A. Thomas and M. D. De Bellis. Pituitary volumes in pediatric maltreatment-related posttraumatic stress disorder. Biol. Psychiatry, 55:752–758, 2004. doi: 10.1016/j.biopsych.2003.11.021.

V. Tseilikman et al. A Rat Model of Post-Traumatic Stress Syndrome Causes Phenotype-Associated Morphological Changes and Hypofunction of the Adrenal Gland. Int. J. Mol. Sci., 22:13235, 2021. doi: 10.3390/ijms222413235.

Y. M. Ulrich-Lai et al. Chronic stress induces adrenal hyperplasia and hypertrophy in a subregion-specific manner. Am. J. Physiol.-Endocrinol. Metab., 291:E965–E973, 2006. doi: 10.1152/ajpendo.00070.2006.

M. van Zuiden et al. Pre-existing high glucocorticoid receptor number predicting development of posttraumatic stress symptoms after military deployment. Am. J. Psychiatry, 168:89–96, 2011. doi: 10.1176/appi.ajp.2010.10050706.

M. van Zuiden et al. Glucocorticoid receptor pathway components predict posttraumatic stress disorder symptom development: a prospective study. Biol. Psychiatry, 71:309–316, 2012. doi: 10.1016/j.biopsych.2011.10.026.

L. Varricchio, E. B. Geer, F. Martelli, M. Mazzarini, A. Funnell, J. J. Bieker, T. Papayannopoulou, and A. R. Migliaccio. Patients with hypercortisolemic Cushing disease possess a distinct class of hematopoietic progenitor cells leading to erythrocytosis. Haematologica, 108(4):1053–1067, 2023. doi: 10.3324/haematol.2021.280542.

A. Verbitsky, D. Dopfel, and N. Zhang. Rodent models of post-traumatic stress disorder: behavioral assessment. Transl. Psychiatry, 10:132, 2020. doi: 10.1038/s41398-020-0806-x.

F. Vogel, L. Braun, S. Zopp, E. Nowak, J. Schreiner, M. Bidlingmaier, M. Reincke, et al. Low-grade inflammation during the glucocorticoid withdrawal phase in patients with Cushing’s syndrome. Eur. J. Endocrinol., 188(4):375–384, 2023. doi: 10.1093/ejendo/lvad041.

K. von Majewski et al. Acute stress responses of autonomous nervous system, HPA axis, and inflammatory system in posttraumatic stress disorder. Transl. Psychiatry, 13(1):36, 2023. doi: 10.1038/s41398-023-02331-7.

K. N. Westlund, G. Aguilera, and G. V. Childs. Quantification of morphological changes in pituitary corticotropes produced by in vivo corticotropin-releasing factor stimulation and adrenalectomy. Endocrinology, 116:439–445, 1985. doi: 10.1210/endo-116-1-439.

R. Yehuda. Status of glucocorticoid alterations in post-traumatic stress disorder. Ann. N. Y. Acad. Sci., 1179:56–69, 2009. doi: 10.1111/j.1749-6632.2009.04979.x.

R. Yehuda, M. T. Lowy, S. M. Southwick, D. Shaffer, and E. L. Giller. Lymphocyte glucocorticoid receptor number in posttraumatic stress disorder. Am. J. Psychiatry, 148:499–504, 1991. doi: 10.1176/ajp.148.4.499.

R. Yehuda, S. M. Southwick, J. H. Krystal, D. Bremner, D. S. Charney, and J. W. Mason. Enhanced suppression of cortisol following dexamethasone administration in posttraumatic stress disorder. Am. J. Psychiatry, 150(1):83–86, 1993. doi: 10.1176/ajp.150.1.83.

R. Yehuda, S. L. Halligan, R. Grossman, J. A. Golier, and C. Wong. The cortisol and glucocorticoid receptor response to low dose dexamethasone administration in aging combat veterans and Holocaust survivors with and without posttraumatic stress disorder. Biol. Psychiatry, 52 (5):393–403, 2002. doi: 10.1016/s0006-3223(02)01357-4.

R. Yehuda, J. A. Golier, S. L. Halligan, M. Meaney, and L. M. Bierer. The ACTH Response to Dexamethasone in PTSD. Am. J. Psychiatry, 161:1397–1403, 2004a. doi: 10.1176/appi.ajp.161.8.1397.

R. Yehuda, J. A. Golier, R.-K. Yang, and L. Tischler. Enhanced sensitivity to glucocorticoids in peripheral mononuclear leukocytes in posttraumatic stress disorder. Biol. Psychiatry, 55(11): 1110–1116, 2004b. doi: 10.1016/j.biopsych.2004.02.010.

R. Yehuda et al. Low urinary cortisol excretion in patients with posttraumatic stress disorder. J. Nerv. Ment. Dis., 178:366–369, 1990. doi: 10.1097/00005053-199006000-00004.

R. Yehuda et al. Influences of maternal and paternal PTSD on epigenetic regulation of the glucocorticoid receptor gene in Holocaust survivor offspring. Am. J. Psychiatry, 171:872–880, 2014. doi: 10.1176/appi.ajp.2014.13121571.

P. R. Zoladz and D. M. Diamond. Current status on behavioral and biological markers of PTSD: a search for clarity in a conflicting literature. Neurosci. Biobehav. Rev., 37:860–895, 2013. doi: 10.1016/j.neubiorev.2013.03.024.

J. V. Zorn et al. Cortisol stress reactivity across psychiatric disorders: A systematic review and meta-analysis. Psychoneuroendocrinology, 77:25–36, 2017. doi: 10.1016/j.psyneuen.2016.11.036.

