## appendix for "Gland-Mass Adaptation Explains the Cortisol Paradox in PTSD"

#### Contents

**Appendix Table S1. Model variables and parameters** 1

**Appendix Table S2. Morning serum cortisol by sex and PTSD status** 2

**Appendix Table S3. Glucocorticoid-action index by window** 2

#### Appendix Table S1. Model variables and parameters

Appendix Table S1. Variables and parameters of the HPA gland-mass model. Hormone rates are in $\text{min}^{-1}$ and gland-mass turnover rates in $\text{day}^{-1}$ (converted to $\text{min}^{-1}$ in the simulations). Because secretion equals degradation for every species, each baseline steady state is normalized to 1, so the steady-state scans use unit rates while the dynamic simulations use the dimensional values. $K_{\text{GR}}$ is the only quantity that differs between PTSD and control. The challenge-specific inputs apply to the named simulation only.

| Symbol | Definition | Scans | Dynamic |
|---|---|---|---|
| *Dynamic variables* | | | |
| $x_1$ | CRH concentration | — | — |
| $x_2$ | ACTH concentration | — | — |
| $x_3$ | Cortisol concentration | — | — |
| $P$ | Pituitary corticotroph functional mass | — | — |
| $A$ | Adrenal cortex functional mass | — | — |
| *Parameters* | | | |
| $b_1$, $a_1$ | CRH secretion, removal | 1 | 0.17 |
| $b_2$, $a_2$ | ACTH secretion, removal | 1 | 0.035 |
| $b_3$, $a_3$ | Cortisol secretion, removal | 1 | 0.06 |
| $b_P$, $a_P$ | Corticotroph growth, turnover | 1 | 0.099 $\text{day}^{-1}$ |
| $b_A$, $a_A$ | Adrenal growth, turnover | 1 | 0.049 $\text{day}^{-1}$ |
| $K_{\text{GR}}$ | GR half-maximal sensitivity | 5 (control) / 2.5 (PTSD) | |
| $n$ | Hill coefficient (GR cooperativity) | 3 | |
| $u$ | Stress input | 1 | 1; TSST 4 pre-stress, 40 pulse |
| *Challenge-specific inputs* | | | |
| $D$ | Dex strength (cortisol-equivalent), Dex–CRH | — | 7 |
| $D_{\text{DST}}$ | Dex strength, suppression test | — | control suppression 72–78% |
| $u_{\text{CRH}}$ | Exogenous CRH, 20-min pulse | — | 0.35 |
| $b_1$, $a_1$ | CRH rates in the Dex–CRH run | — | 0.017 |

### Appendix Table S2. Morning serum cortisol by sex and PTSD status

Appendix Table S2. Crude and age-standardized morning serum cortisol (nmol/L) by sex and PTSD status, adults aged 18–70. Standardization uses the combined PTSD-plus-control unique-individual age distribution as the common reference, so the standardized means differ only by the PTSD effect and not by age structure. Standardized contrasts: men $\Delta = -19.8$ nmol/L ($-4.6\%$), $g = -0.11$ (95% CI $-0.20$ to $-0.03$), $p = 0.012$; women $\Delta = -20.1$ nmol/L ($-4.5\%$), $g = -0.11$ ($-0.17$ to $-0.04$), $p = 0.001$. Unique individuals are counted once each across age bins.

| Sex | Group | $n$ (tests) | $n$ (indiv.) | Crude mean ± SD | Age-std. mean ± SE |
|---|---|---|---|---|---|
| Men | Control | 49,085 | 38,085 | $433 \pm 172$ | $432.4 \pm 0.8$ |
| | PTSD | 505 | 404 | $413 \pm 168$ | $412.6 \pm 7.8$ |
| Women | Control | 129,995 | 97,814 | $444 \pm 185$ | $443.4 \pm 0.5$ |
| | PTSD | 904 | 730 | $419 \pm 175$ | $423.3 \pm 6.1$ |

### Appendix Table S3. Glucocorticoid-action index by window

Appendix Table S3. Glucocorticoid-action index, in SD units of the control distribution. Person-level: each person contributes one value per window. $p$ from 2,000 permutations of arm labels within sex; 0.0005 is the resolution limit. PTSD and control columns give the number of persons.

| Window | $n$ PTSD | $n$ control | PTSD − control | 95% CI | $g$ |
|---|---|---|---|---|---|
| Baseline −10 to −6 y | 30,528 | 217,097 | +0.0375 | [+0.0332, +0.0417] | +0.105 |
| Prodrome −6 to −1 y | 35,555 | 245,586 | +0.0491 | [+0.0451, +0.0530] | +0.137 |
| Diagnostic −1 to +0.5 y | 31,711 | 178,654 | +0.0584 | [+0.0538, +0.0630] | +0.150 |
| Early +0.5 to +3 y | 25,495 | 167,820 | +0.0538 | [+0.0488, +0.0588] | +0.140 |
| Late +3 to +10 y | 16,447 | 116,469 | +0.0537 | [+0.0474, +0.0599] | +0.143 |
